# Characterization of Mayer wave oscillations in functional near-infrared spectroscopy using a physiologically informed model of the neural power spectra

**DOI:** 10.1101/2021.09.01.458637

**Authors:** Robert Luke, Maureen J Shader, David McAlpine

## Abstract

**Significance:** Mayer waves are spontaneous oscillations in arterial blood pressure that can mask cortical hemodynamic responses associated with neural activity of interest.

**Aim:** To characterize the properties of oscillations in the fNIRS signal generated by Mayer waves in a large sample of fNIRS recordings. Further, we aim to determine the impact of short-channel correction for the attenuation of these unwanted signal components.

**Approach:** Mayer wave oscillation parameters were extracted from 310 fNIRS measurements using the Fitting Oscillations & One-Over-F (FOOOF) method to compute normative values. The effect of short-channel correction on Mayer wave oscillation power was quantified on 222 measurements. The practical benefit of the short-channel correction approach for reducing Mayer waves and improving response detection was also evaluated on a subgroup of 17 fNIRS measurements collected during a passive auditory speech detection experiment.

**Results:** Mayer-wave oscillations had a mean frequency of 0.108 Hz, bandwidth of 0.04 Hz, and power of 3.5 μM^2^/Hz. The distribution of oscillation signal power was positively skewed, with some measurements containing large Mayer waves. Short-channel correction significantly reduced the amplitude of these undesired signals; greater attenuation was observed for measurements containing larger Mayer-wave oscillations.

**Conclusions:** A robust method for quantifying Mayer-wave oscillations in the fNIRS signal spectrum was presented and used to provide normative parameterization. Short-channel correction is recommended as an approach for attenuating Mayer waves, particularly in participants with large oscillations.

## 1. Introduction

The ability to explore and understand the brain requires the differentiation of signals that arise from the neural activity of interest from signals that are generated by other factors or non-neural processes. Neuroimaging techniques that estimate the concentration of oxygenated and/or deoxygenated hemoglobin within the cortical blood flow, including functional magnetic resonance imaging (fMRI) and function near-infrared spectroscopy (fNIRS), are susceptible to disruptions from non-neural cardiovascular oscillations in the signal (e.g., Tong et al., 2012). For example, oscillations corresponding to heart rate (~1 Hz) and respiratory rate (~0.3 Hz) are present within the broad frequency spectrum of the measured signal and can be removed or reduced through filtering. Another cardiovascular component is Mayer waves, which are spontaneous oscillations in arterial blood pressure with a frequency of ~0.1 Hz (Ghali and Ghali, 2020; Julien, 2006). Mayer waves are not easily removed from hemodynamic signatures of brain activity as they tend to occur on a time course often confounded with the frequency of a sensory task, for example, and/or the cortical hemodynamic response to that task.

Assuming Mayer waves are predominately manifested as oscillations in oxygenated hemoglobin within the scalp (Zhang et al., 2015), Yücel et al. (2016) inserted synthetic hemodynamic response functions (HRFs) into the resting-state recordings from 17 participants in an fNIRS study to demonstrate that large Mayer waves significantly reduce the accuracy of the estimated synthetic hemodynamic response. Further, they demonstrated that utilizing dedicated source-detector pairs for measuring predominantly extra-cerebral activity (8 mm source-detector spacing), referred to as short-channels, reduced the impact of Mayer waves on the estimation of the cortical hemodynamic response, and that this effect was more pronounced when the true change in the concentration of oxygenated hemoglobin was small relative to the magnitude of the Mayer wave. This confirms that short-channel correction improves the accuracy with which HRFs are estimated for synthetic cortical responses when large Mayer waves are present. However, the effect of short-channel correction on the characteristics of Mayer waves and the effect of those oscillations on the detection of HRFs in real measurements with event-driven responses has yet to be quantified.

Although many techniques have been proposed to reduce or remove the contribution from Mayer waves to cortical hemodynamic responses (e.g., von Lühmann et al., 2020), it is commonly reported that for a small proportion of subjects, HRFs cannot be recovered due to the presence of large-amplitude Mayer waves leading to these participants being excluded from group-level analyses of hemodynamic responses (Yücel et al., 2016). A better understanding of how Mayer waves might be quantified will facilitate the development of objective criteria for rejecting hemodynamic responses contaminated by Mayer waves, and in developing and evaluating algorithms to reduce systemic contributions to hemodynamic responses in neuroimaging data.

Mayer wave activity in fNIRS measurements is typically quantified by aggregating the signal activity within a frequency band centered at 0.1 Hz and a bandwidth of 0.08 Hz (e.g., 0.06-0.14 Hz), although the exact frequency band varies between studies (Kirilina et al., 2013; Pinti et al., 2019; Yücel et al., 2016; Zhang et al., 2015). This approach is commonplace across neuroimaging modalities, including EEG, ECOG, and MEG (de Beeck and Nakatani, 2019). However, recent reports suggest that using predefined frequency bands can generate incorrect estimates of oscillatory power, especially if the center frequency of oscillatory activity is intrinsically variable (Lansbergen et al., 2011). Further, using predefined frequency bands conflates changes in the power of oscillations of interest with shifts in center frequency, changes in broadband power, and changes in the non-oscillatory components of neuroimaging signals (Donoghue et al., 2020a). To overcome these shortcomings and facilitate accurate and objective parameterization of oscillatory activity in neuroimaging data, Donoghue et al. (2020b) suggested analyzing oscillatory activity by modelling the aperiodic spectral features— decreasing power across increasing frequencies—as well as the periodic/oscillatory components. Rather than ignoring or correcting for these components, which ignores their possible physiological correlates, the algorithm employed by Donoghue et al. (2020b) provides a physiologically informed model of the neural power spectra by modeling both distinct functional processes. This approach, called “Fitting Oscillations & One-Over-F” (FOOOF) and provided by the authors as open-source software, allows for the identification of oscillatory activity in hemodynamic signals without the requirement of predefined and specific frequency bands of interest. It is, therefore, well suited to identifying and parameterizing Mayer wave activity in fNIRS measurements.

Here, we quantify the typical frequency and power distribution of Mayer waves from over 300 fNIRS recordings using the FOOOF algorithm. This accords with the approach of Yücel et al. (2016), who utilized synthetic data to demonstrate the ground truth of the effect of short-channel correction as a function of the amplitude of HRFs and Mayer waves. The mitigation effect of using short-separation channel regression for reducing contamination of hemodynamic responses by Mayer waves is evaluated on over 200 fNIRS recordings. In addition, the impact of Mayer waves on the detection of auditory-evoked hemodynamic responses, which typically comprise small responses and poor signal-to-noise ratios (Luke et al., 2021), is evaluated in a subgroup of recordings from 17 participants.

## 2. Methods

### 2.1 Data

The data used in this study were aggregated from a variety of research studies conducted between 2018 and 2021. Measurements were obtained under the Macquarie University Ethics Application References 52020640814625 and 5201500948. In total, 310 measurements are included in this dataset. The studies contained a variety of optode-placement montages, sample rates, experimental designs, tasks, and stimuli. As such, it is not appropriate to compare task-evoked responses across measurements. However, a comparison of systemic oscillations within the signal, which are unrelated to the task-evoked aspect of the data (e.g., Mayer waves), is valid.

All participants were seated in a sound-attenuating booth in a comfortable chair for the duration of the experiments. NIRS data were recorded using a continuous-wave NIRx NIRScoutX device with APD detectors. Measurements were obtained with between 8-16 sources and detectors. Experiments lasted between 15 and 80 minutes. All data were stored in BIDS data format (Gorgolewski et al., 2016). Data were processed with MNE-Python (Gramfort et al., 2013; Gramfort et al., 2014), MNE-NIRS (Luke et al., 2021), and Nilearn (Abraham et al., 2014; Thirion et al., 2021).

### 2.2 Quantification of Mayer-wave parameters

The FOOOF method of Donoghue et al. (2020b) was employed to quantify oscillations in the fNIRS signal associated with Mayer waves. This method was designed to extract oscillations in neural signals while accounting for the natural structure of the frequency spectrum of neuroimaging signals. As such, it is well suited for quantifying Mayer waves; the method extracts peaks in the power spectra without the need to predefine analysis parameters, such as the signal center frequency or bandwidth.

To quantify the parameters of Mayer waves, a minimal processing pipeline was applied to covert the raw data to oxyhemoglobin concentration. Specifically, the raw data were first converted to optical density. Channels with a scalp coupling index less than 0.7 were excluded from further analysis (Pollonini et al., 2014), and the data were down sampled to 1.7 Hz, to match the lowest sampling rate of any measurement in the dataset. The data were then converted to hemoglobin concentration using the modified Beer-Lambert law with a differential pathlength factor of 0.1. Channels with a source-detector distance of less than 1.5 or more than 4.5 cm were excluded from further analysis. Finally, as the oxyhemoglobin measurement contains a greater contribution from Mayer wave signals than deoxyhemoglobin (Zhang et al., 2015), only the oxyhemoglobin signal was retained for further analysis.

The power spectral-density was then calculated for each channel using the Welch method with a Hamming window comprising 300 samples and 150 samples of overlap, and the spectra averaged across all channels per measurement to generate a single power spectral-density estimate per recording (in units of μM^2^/Hz). These spectra were then passed to the FOOOF software. The FOOOF software returns estimates of the aperiodic component and oscillations in the signal; if multiple oscillations are detected (e.g., respiratory rate) then the oscillation closest to 0.1 Hz is retained for further analysis as it is assumed to contain a Mayer-wave component. The output of this analysis is a single estimate of the oscillations present in a recording. Utilizing the software provided by the FOOOF authors is recommended, as it reports the oscillations within the likely Mayer wave range in terms of their center frequency in Hz, bandwidth in Hz, and log power in arbitrary units.

### 2.3 Effect of short-channel correction

We evaluated a subset of 222 files that contained data from short-separation channels to quantify the effect of systemic signal attenuation on the presence of Mayer-wave oscillations in the fNIRS signal. After estimating Mayer-wave parameters when no attenuation was applied to the systemic signal (see Section 2.2), we employed the same approach with the additional step of reducing the systemic component of the hemodynamic response through the assessment of activity in short-channels being applied after resampling. Many methods have been proposed for attenuating systemic components. This study utilized the method of short-channel regression based on the nearest short channel (algorithm as provided by NIRx) (Saager and Berger, 2005; Scholkmann et al., 2014). As a result, the parameters of Mayer-wave oscillations (center frequency, bandwidth, and power) were estimated with and without short-channel based correction of systemic signals in the hemodynamic response.

A Bland-Altman analysis (Altman and Bland, 1983) was used to determine whether short-channel correction affected Mayer waves differentially as a function of their amplitude. The bootstrapping approach of Ho et al. (2019) was used to compare the magnitude of Mayer-wave oscillations with and without short-channel correction.

### 2.4 Single dataset example

To demonstrate the practical utility of measuring the parameters of Mayer waves, a single, publicly available dataset of 17 participants was analyzed (Luke et al., 2021). The presence of an evoked response to auditory-speech stimuli was quantified at an individual level to determine whether data from participants with larger Mayer waves benefited from correction of systemic components in their hemodynamic responses. Mayer waves were quantified for the data from each participant as described in Section 2.2.

The data were analyzed using a generalized linear model (GLM), as, due to the statistical properties of the noise in fNIRS signals (Huppert, 2016), this is suggested to be more appropriate than averaging the data. The data were analyzed as described in Luke et al. (2021). Briefly, the raw data were first converted to optical density, and then to hemoglobin concentration using the modified Beer-Lambert Law with a differential pathlength factor of 0.1. They were then down sampled to 0.6 Hz, and a GLM was fitted to all channels using a 3-sec boxcar function convolved with a glover model. A cosine drift was included with components up to 0.01 Hz. Two analyses were performed, one including the mean of the short channels as a regressor, and one without any short-channel regressor. Channels over the left and right superior temporal gyri, comprising typical auditory regions of interest, were then combined into a single region of interest using a weighted average of the GLM estimates (Shader et al., 2021). If the p-value of the regressor component of the speech task in the GLM estimate was less than 0.05 (i.e., a significant HbO response evoked by auditory speech) a response was deemed present. As such, for each participant, a task-locked neural response was quantified as either present or absent for both analyses, with and without short-channel regression.

## 3. Results & Discussion

Accurate estimation of neuroimaging signal components is essential for precise measurements of brain activity. Here, we investigate the use of the FOOOF method for characterizing Mayer-wave oscillations in the fNIRS signal. The FOOOF method estimates oscillatory activity in the neuroimaging measurement that is not a component generated by the aperiodic structure of the signal. Thus, characterization of the peaks in the power spectrum are possible without requiring predefined bandwidths. Figure 1 illustrates several representative fits provided by the FOOOF algorithm applied to fNIRS data. The algorithm appropriately models the data in the presence of small (Figure 1a) and large (Figure 1b) oscillation peaks, for oscillations not centered exactly at 0.1 Hz (Figure 1b), and in the presence of additional oscillations in the data (likely respiratory rate) (Figure 1c).

**Figure 1:**
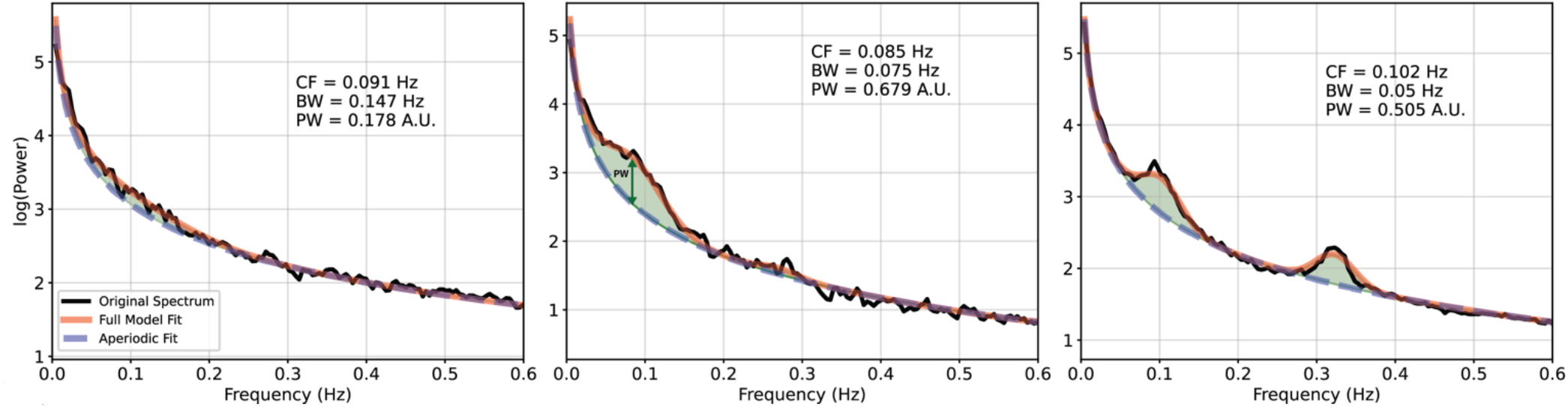
Examples of FOOOF fit of fNIRS data. The black line represents the power spectral density of the signal, the red line represents the complete model fit of the FOOOF algorithm, with the blue dashed line indicating the aperiodic portion of the signal, and green shaded regions marking oscillations in the signal as peaks rising above the aperiodic component. Three examples are provided to demonstrate the appropriateness of the algorithm to fNIRS data in different situations: a) provides an example of a measurement with very small Mayer wave oscillation, b) demonstrates an oscillation that is not centred at the expected 0.1 Hz frequency, and c) demonstrates a measurement with additional substantial oscillatory activity at 0.32 Hz which likely represents the breathing/respiratory rate. CF = center frequency (Hz). PW = power estimation (A.U.). BW = bandwidth (Hz).

The underlying code to run the FOOOF algorithm is available at the original authors’ website (Donoghue et al., 2020b), and a convenient wrapper to run the analysis with fNIRS data is provided as part of the MNE-NIRS package (Luke et al., 2021).

### 3.1 Quantification of Mayer-wave parameters on a cohort of 310 measurements

The FOOOF algorithm was applied to a cohort of 310 measurements. An oscillatory component was detected in all but 4 measurements. The mean center frequency of the oscillations was 0.108 Hz and the median value was 0.097 Hz (see Figure 2a). This was expected as the analysis approach actively selected the frequency component closest to 0.1 Hz in instances when more than one oscillatory peak was identified in the signal. To confirm that the selected peaks were indeed related to Mayer-wave activity, the model was rerun while retaining oscillations with frequencies closest to typical cutoff frequencies used in frequency-band aggregates of Mayer waves, including 0.05 and 0.15 Hz. With these modified parameters, the median frequency of the resulting oscillatory component was essentially unchanged at 0.105 and 0.109 Hz for the lower and upper cutoff frequencies, respectively, demonstrating that the model was fitting robust oscillatory components in the data related to Mayer-wave activity.

**Figure 2:**
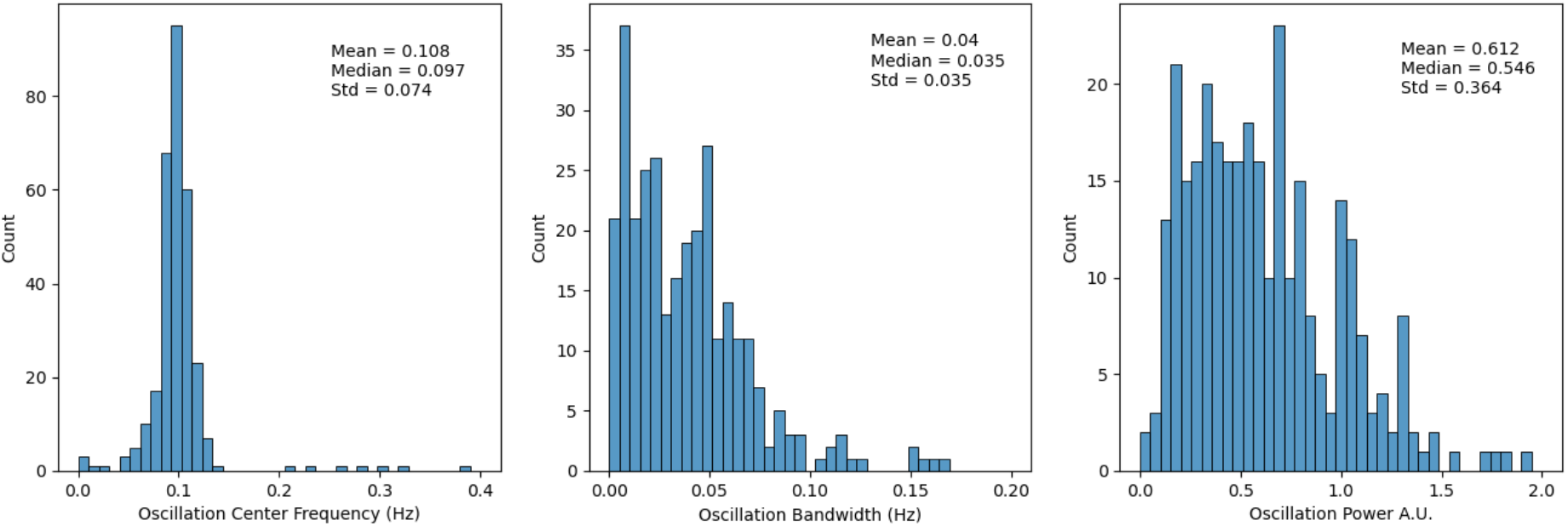
Quantification of Mayer-wave parameters. a) center frequency of oscillation components, b) oscillation bandwidth, c) power of oscillation component. Note that the power of Mayer waves is not normally distributed, with some participants having much larger values than the group majority.

The bandwidth of the oscillations was highly non-normally distributed with a median value of 0.035 Hz (Figure 2b), corresponding to an average Mayer-wave frequency range of approximately 0.065-0.135 Hz. Similarly, Mayer-wave power was not uniformly distributed across participants (Figure 2c). Instead, the median log power of oscillations was 0.55 A.U., and the distribution was positively skewed. This corresponds to an oscillatory component in the fNIRS signal of 3.5 μM^2^/Hz. Nineteen of 310 measurements (6%) contained Mayer waves that were 2 standard deviations above the median oscillation power, largely consistent with anecdotal reports of a small proportion of participants having particularly large Mayer waves. Unbiased quantification of the parameters of these oscillations while including the aperiodic component in the spectra will assist in the development of algorithms for identifying and attenuating these unwanted signals in hemodynamic responses.

### 3.2 Effect of short-channel correction on Mayer waves

Next, the degree to which the power of Mayer waves is attenuated following short-channel correction was evaluated, and it was assessed if any characteristic of the uncorrected Mayer-wave oscillations (i.e., greater oscillatory power) was associated with larger power attenuation after correction. Of the 310 recordings in the dataset, 222 contained short-channel data, providing a means by which to assess the effect of short-channel correction on Mayer-wave power. These measurements were reanalyzed to quantify the center frequency, bandwidth, and power of the Mayer waves after short-channel systemic correction was applied to the long-channel data.

Applying short-channel correction did not significantly alter the frequency distribution or bandwidth of Mayer-wave components in the signal (Figure 3a). A small but significant reduction in the signal power of Mayer waves was apparent when short-channel systemic correction was applied to the entire dataset (effect=−0.0459, 95%CI [−0.066, −0.027], N=222, *p*<0.001).

**Figure 3:**
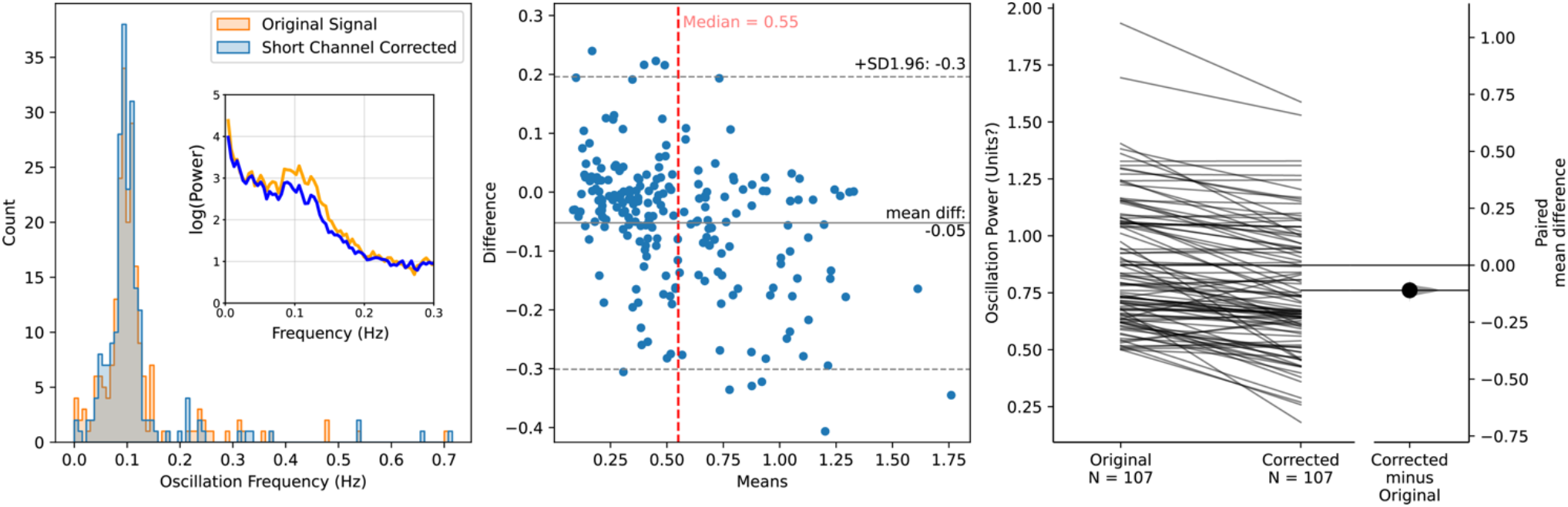
Effect of short-channel correction on the power of Mayer-wave oscillations a) Distribution of frequency component of Mayer waves with and without short channel-correction applied. Inset illustrates an example measurement with and without short-channel correction, note the increase in power around 0.1 Hz is reduced when correction is applied, b) Bland-Altman plot illustrating the difference in Mayer-wave power when short-channel correction is applied as a factor of the oscillation power, c) paired comparison illustrating the effect of short-channel correction on oscillation power. Note that short-channel correction does not affect the frequency of the oscillation component but does reduce the power of the oscillation, particularly for measurements containing a large Mayer wave oscillatory component.

The reduction in Mayer-wave power was more pronounced in measurements with a large systemic component in the uncorrected data. Bland-Altman analysis (Figure 3b) demonstrates that the difference in oscillation power when the correction is applied varies as a function of oscillation power. As such, we analyzed the measurements with the largest Mayer-wave power using a median split to divide the dataset into two groups. The group with larger Mayer waves (107 measurements) showed a reduction of 0.11 in log power when short-channel correction was applied (effect=−0.11, 95%CI [−0.136, −0.086], N=107, *p*<.001), whereas the group with smaller Mayer-wave oscillations showed no consistent change in oscillation power following short-channel correction (effect=−0.0018, 95%CI [−0.0145, 0.021], N=115, *p*=.841). On this basis, short-channel correction to reduce Mayer-wave components in hemodynamic signals provides significant benefit when Mayer-wave components are relatively large, supporting the HRF modeling of Yücel et al. (2016). Further, short-channel correction does not systematically alter the oscillation component when applied to measurements with small Mayer waves, indicating that short-channel systemic component correction can be broadly applied to fNIRS measurements without unwanted signal distortion.

### 3.3 The effect of short-channel correction on the amplitude of evoked responses

Systemic noise can mask the presence of neural responses in fNIRS measurements. Accounting for data obtained from short channels in the signal processing pipeline may mitigate unwanted systemic components and improve the detection of neural responses. As such, we analyzed the measurements from a publicly available dataset (Luke et al., 2021) comprising fNIRS data of 17 participants, to determine if measurements with large Mayer waves particularly benefited from short-channel correction.

As for the larger data set (Section 3.2), data were divided into two groups using a median split of the oscillation power in Mayer waves. Without short-channel regression, just two of nine measurements with the largest Mayer waves were observed to contain significant evoked responses to auditory-speech stimuli. When short-channel regression was employed, however, this increased to eight of nine measurements, suggesting that short-channel correction does indeed improve the detection of auditory-evoked cortical responses in participants with large Mayer waves. This data was collected with an experimental design with randomized inter-stimulus-interval times to minimize listener expectation and mitigate the effect of Mayer wave contributions. Even greater benefits from short-channel correction may be observed for data collected without randomized inter-stimulus-intervals.

## 4. Conclusion

A method to quantify the presence of Mayer-wave oscillations in fNIRS measurements was introduced based on the well-established and publicly available algorithm of Donoghue et al. (2020b). The technique was applied to a dataset of 310 fNIRS measurements and the population normative values are reported. The distribution of oscillation power in Mayer waves was found to be positively skewed. From a subset of 222 recordings, it was demonstrated that applying short-channel correction significantly reduces contamination by Mayer-wave oscillations in responses containing large oscillatory activity, but does not consistently modify the oscillation properties of responses with smaller Mayer waves. The applicability of this technique for quantifying Mayer-wave oscillations in an evoked-response experiment was demonstrated on a publicly available dataset. From this, we recommend using the FOOOF method to quantify Mayer-wave oscillatory activity and to evaluate the efficacy of algorithms designed to mitigate systemic components in hemodynamic responses, including through the use of short-channel based algorithms when assessing hemodynamic responses generated using fNIRS.

## 5. Code, Data, & Materials Availability

The FOOOF method is available at Donoghue et al. (2020b) website [https://fooof-tools.github.io/fooof/]. An implementation of the algorithm for use with fNIRS data will be made available at the MNE-NIRS website.

## 6. Acknowledgments/Funding Sources

This study was supported by an Australian Research Council Laureate Fellowship (No. FL160100108) awarded to David McAlpine.

## Notes

### Competing Interest Statement

The authors have declared no competing interest.

### Summary of Updates

Figure 2 revised

